# Mitoregulin self-associates to form likely homo-oligomeric pore-like structures

**DOI:** 10.1101/2024.07.10.601956

**Authors:** Connor R. Linzer, Colleen S. Stein, Nathan H. Witmer, Zhen Xu, Nicholas J. Schnicker, Ryan L. Boudreau

**Affiliations:** Department of Internal Medicine, Carver College of Medicine, University of Iowa, Iowa City, IA, United States; Molecular Medicine Graduate Program, Carver College of Medicine, University of Iowa, Iowa City, IA, United States; Protein and Crystallography Facility, University of Iowa Carver College of Medicine, Iowa City, IA, United States; Department of Molecular Physiology and Biophysics, University of Iowa Carver College of Medicine, Iowa City, iA, United States; Fraternal Order of Eagles Diabetes Research Center and Abboud Cardiovascular Research Center, Carver College of Medicine, University of Iowa, Iowa City, IA, United States

**Author notes:** To whom correspondence should be addressed: Ryan L. Boudreau 4334 PBDB University of Iowa Department of Internal Medicine Iowa City, IA 52242.

## Abstract

We and others previously found that a misannotated long noncoding RNA encodes for a conserved mitochondrial transmembrane microprotein named Mitoregulin (Mtln). Beyond an established role for Mtln in lipid metabolism, Mtln has also been shown to more broadly influence mitochondria, boosting respiratory efficiency and Ca^2+^ retention capacity, while lowering ROS, yet the underlying mechanisms remain unresolved. Prior studies have identified possible Mtln protein interaction partners; however, a lack of consensus persists, and no claims have been made about Mtln’s structure. We previously noted two key published observations that seemingly remained overlooked: 1) endogenous Mtln co-immunoprecipitates with epitope-tagged Mtln at high efficiency, and 2) Mtln primarily exists in a ∼66 kDa complex. To investigate if Mtln may self-oligomerize into higher-order complexes, we performed co-immunoprecipitation, protein modeling simulations, and native gel assessments of Mtln-containing complexes in cells and tissues, as well as tested whether synthetic Mtln protein itself forms oligomeric complexes. Our combined results provide strong support that Mtln self-associates and likely forms a hexameric pore-like structure.

## INTRODUCTION

Less than 5% of our genomic landscape encodes for proteins, yet studies have found that much of our DNA gives rise to thousands of uncharacterized RNAs, including long non-coding RNAs (lncRNAs), RNAs >200 nucleotides long that are thought to lack protein-coding open-reading frames (ORFs). However, newer reports suggest that many lncRNAs may be misannotated and actually encode microproteins (<100 amino acids, below the original cut-off for defining ORFs)^1^. Our group and others have discovered an array of conserved microproteins that are diverse in nature and play broad cellular roles, e.g. in Ca^2+^ pump modulation^2,3^, mRNA processing^4^, and mitochondrial metabolism^5,6^. Given the sheer scale and diversity of microproteins, further research is necessary to continue to fully elucidate their potential molecular actions and broader physiologic impacts.

We previously discovered that the lncRNA LINC00116 is misannotated and harbors a highly-conserved (e.g. fish to humans) small ORF that gives rise to a 56-amino acid mitochondrial transmembrane microprotein that we named Mitoregulin (Mtln)^5^. We detailed Mtln’s strong binding to cardiolipin and broad influence on multiple mitochondrial properties and functions, including membrane potential, oxidative phosphorylation (OXPHOS), reactive oxygen species (ROS), calcium handling, and lipid metabolism. To date, seven independent groups have claimed co-discovery of Mtln^5,7–12^, and there is reasonably strong consensus on its influence on fatty acid (beta-) oxidation, as well as several consistent data points across papers supporting its broader impacts on mitochondrial OXPHOS, ROS and membrane potential. However, the precise molecular function of Mtln remains unclear, and there are discrepancies regarding its localization (e.g. inner versus outer mitochondrial membranes^1,5,9,12^) and potential protein binding partners. Overall, protein interaction studies (e.g. via co-immunoprecipitation coupled with mass spectrometry), typically done in the setting of forced overexpression of epitope-tagged Mtln constructs, have yielded mostly non-reproducible and/or unconvincing data on Mtln interactors, several of which have been proposed (e.g. Hadha/b, Cyb5r3, Atp5a, and Ndufa7). Along these lines, we previously reported SDS-PAGE and blue native PAGE (BN-PAGE) western blot data for Mtln showing multiple bands and a broad/smeared pattern, which we interpreted as Mtln potentially being a “sticky” protein that associates with many mitochondrial complexes, perhaps serving as an assembly factor^5^. However, Zhang et al. recently reported that Mtln tends to aggregate upon sample freezing, leading to the artifactual appearance of high molecular weight bands and smearing on BN-PAGE^12^. Thus, it is possible that our published findings may have resulted from technical artifact, given that our previous samples were indeed frozen prior to mitochondrial isolation and BN-PAGE. In fresh (i.e. not previously frozen) samples, Zhang et detected non-aggregated Mtln primarily in the lower molecular weight range, migrating as an intense band at ∼66 kDa. Notably, we also observed this prominent ∼66 kDa Mtln-containing complex in our BN-PAGE studies, and this captured our interest^5^. To date, this complex remains of unknown composition, and none of the proposed Mtln interacting proteins identified by mass spectrometry appear to co-migrate at this position. Together, these observations have driven us to consider the possibility that Mtln’s strongest protein binding partner might be itself. With this view, we noticed published data showing that native Mtln efficiently co-immunoprecipitated with epitope-tagged Mtln constructs (Figure 6B in Xiao et al.^9^), and this has seemingly been overlooked. Thus, it is plausible that the ∼66 kDa band may correspond to a homo-oligomeric complex of Mtln subunits. Indeed, homo-oligomerization has been reported for some microproteins, including those that bind to and regulate sarco(endo)plasmic reticulum calcium ATPases^13^, as well as PIGBOS, a mitochondrial microprotein that regulates endoplasmic reticulum stress responses^14^.

To further investigate whether Mtln may self-oligomerize, we performed our own co-immunoprecipitation studies using epitope-tagged Mtln constructs, ran *in silico* protein modeling and oligomerization simulations, assessed migration of complexes containing native or epitope-tagged Mtln by BN-PAGE, and tested whether synthetic Mtln protein can oligomerize to form higher-order complexes. Our combined results provide strong support that Mtln is a self-associating microprotein and reveal the intriguing possibility that Mtln oligomerizes to form a hexameric pore-like structure.

## RESULTS

### Native Mtln co-immunoprecipitates with epitope-tagged Mtln

We previously noted that, in a report by Xiao et al.^9^, a prominent band corresponding to endogenous Mtln was detected by western blot performed on samples after immunoprecipitation of FLAG epitope-tagged Mtln, suggesting that Mtln likely self-associates in some manner. However, the authors did not mention this and instead focused on possible Mtln interaction with Ndufa7, which in comparison to native Mtln, appeared to be much more weakly co-immunoprecipitated (based on input:pulldown ratios). To further examine this, we generated a series of plasmids expressing V5 epitope-tagged (N-terminal or internal) human Mtln constructs to determine if untagged native Mtln is also pulled down with anti-V5 immunoprecipitation (**Fig. 1A**). Additionally, we generated Mtln knockout (KO) A549 cells (a human lung adenocarcinoma cell line that expresses high levels of Mtln) using CRISPR-Cas9 technology (**Fig. 1B**). V5-epitope tagged Mtln constructs were first transfected into Mtln-KO cells to confirm mitochondrial localization prior to immunoprecipitation studies; anti-Mtln immunolabeling overlapped MitoTracker^TM^ staining and was absent in non-transfected cells (**Fig. S1**). These plasmids were subsequently transfected into wild-type (WT) A549 cells alongside an empty vector control, and cell lysates were subjected to anti-V5 immunoprecipitation and subsequent anti-Mtln western blot analysis (**Fig. 1C**). Endogenous Mtln (migrating at ∼11 kDa) was evenly expressed across input samples and appeared to be co-immunoprecipitated along with Mtln constructs tagged with V5 on the N-terminus (nV5, ∼13 kDa) but not with empty vector control nor Mtln constructs harboring internal V5 tag (intV5), the latter of which may hinder association with native Mtln and instead provides a control indicating that Mtln does not simply interact with V5 tags.

**Figure 1:**
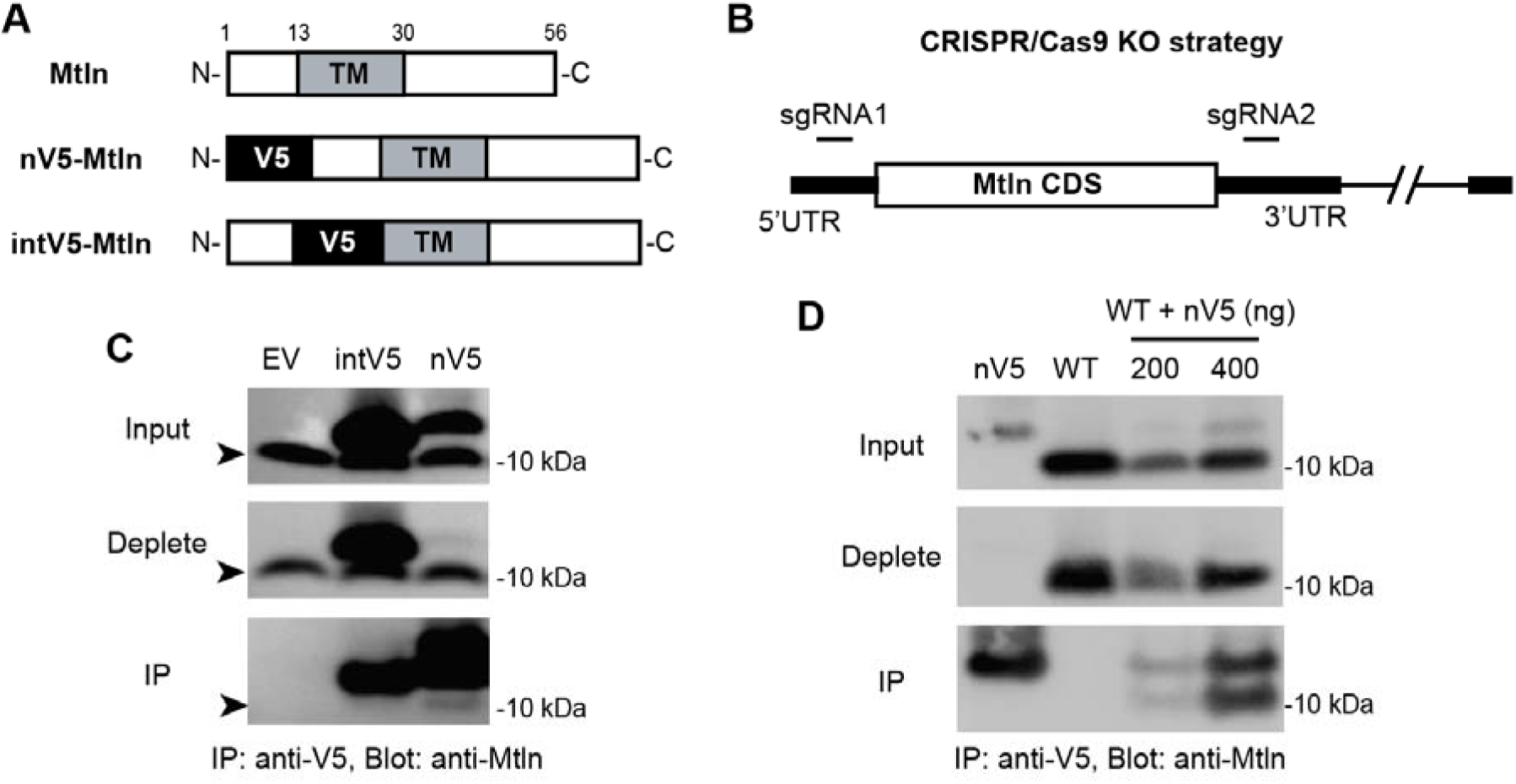
Native Mtln co-immunoprecipitates with epitope-tagged Mtln. **A.** Schematic of plasmid-based human Mtln expression constructs used in immunoprecipitation experiments. V5 tag is represented in relation to the N-terminus (N), C-terminus (C), and transmembrane domain (TM) of Mtln, which is 56 amino acids in length. **B.** Overview of CRISPR/Cas9 knockout strategy used to generate A549 Mtln knockout (Mtln-KO) cells. **C.** An empty vector (EV) control, internal V5 tagged Mtln (intV5), and nV5 tagged Mtln (nV5) were transfected into wildtype A549 cells, and lysates were subsequently subjected to anti-V5 immunoprecipitation (IP). The input, deplete, and IP fractions were assessed by western blot using an anti-Mtln antibody; arrowheads point to lower bands corresponding to endogenous Mtln and upper bands correspond to V5-tagged Mtln. **D.** Wildtype Mtln (WT) and nV5 tagged Mtln (nV5) were co-transfected into Mtln-KO cells and subjected to anti-V5 IP. The input, deplete, and IP fractions were assessed by western blot using an anti-Mtln antibody.

We noted that the amount of pulled down nV5-Mtln far exceeded that of co-immunoprecipitated endogenous Mtln and speculated that endogenous Mtln may already be stably interacting with itself (or other proteins) prior to nV5-Mtln transgene expression. To circumvent this potential caveat, we used a native (untagged) Mtln plasmid construct to allow for simultaneous co-expression with V5-tagged Mtln in Mtln-KO cells. Plasmids expressing native Mtln and nV5-Mtln were transfected individually or together into Mtln-KO cells, and cell lysates were once again subjected to anti-V5 immunoprecipitation and anti-Mtln western blot analysis (**Fig. 1D**). Notably, native Mtln was better expressed compared to nV5-Mtln; however, anti-V5 immunoprecipitation clearly pulled down and enriched nV5-Mtln, while co-precipitating a near equal amount of native Mtln. One interpretation of these findings is that the majority of nV5-Mtln proteins were bound to native Mtln proteins, suggesting a high degree of interaction. To assess whether these observations could be artifactual and reflective of generic pulldown of mitochondrial membranes and associated proteins, we blotted our eluted samples using an OXPHOS cocktail antibody and found no evidence for respiratory complex subunits being co-immunoprecipitated with nV5-Mtln (**Fig. S2**), supporting the specificity of nV5-Mtln interaction with WT Mtln in these experiments. Overall, these data provide further evidence supporting that Mtln self-associates.

### Simulation of Mtln self-association predicts stable hexamer formation

To begin examining the potential nature of this self-association, we first generated simulated models of homo-oligomeric structures *in silico*. We obtained singlet microprotein structural models for human Mtln using AlphaFold^15^ and i-Tasser^16^ on default settings, with each showing a predominant alpha-helical protein structure (**Fig. 2A**). To determine the number of Mtln subunits that could form the most probable homo-oligomeric complex, we input these structural models into the GalaxyHomomer web-based tool^17^. Models of 3-12 Mtln subunits were analyzed for each of the singlet structures (one for AlphaFold and the top two for i-Tasser), and the relative docking scores and normalized interface areas were obtained for each multimer. Of note, higher docking scores and interface areas (per subunit) indicate increased probability of forming more stable (i.e. likely) homo-oligomeric structures. Homomers of only two subunits were not considered, as only antiparallel models with misaligned transmembrane domains were generated. For the remaining multimers, we only considered models showing aligned and crudely parallel transmembrane domains, given considerations for membrane constraints (alignment and width) that GalaxyHomomer does not consider for its modeling. Across all possible multimeric complexes that were examined, the i-Tasser top-ranked singlet model produced a hexameric complex with the highest docking score (605.13) and interface area per subunit (1271.02). In addition, a combined analysis of the measurements for both parameters across the three singlet models provides overall support that Mtln homo-oligomerization is predicted to most likely produce a hexameric complex (**Fig. 2B**). Visual assessment of the predicted hexameric structure revealed an intriguing pore-like complex with a cluster of positively-charged residues that may project into the mitochondrial intermembrane space from the inner mitochondrial membrane and/or into the cytosol from the outer mitochondrial membrane (**Fig. 2C**). Importantly, these hexameric complexes also presented with plausible membrane spanning widths (∼2.8 nM, close to known widths of mitochondrial membrane lipid bilayers), which grew narrower in complexes with 7 to 10 subunits, as these had more severely angled alpha-helices (**Fig S3**). Interestingly, 11-mer and 12-mer complexes also had membrane-spanning regions of adequate widths (∼3 nM), which is notable since we have also observed the presence of ∼120-140 kDa Mtln-containing complexes (Stein et al.^5^ and see below). Overall, these predictive data provide unbiased support for the possibility that Mtln may self-oligomerize to form various pore-like structures, while identifying the hexameric form as a strong candidate.

**Figure 2:**
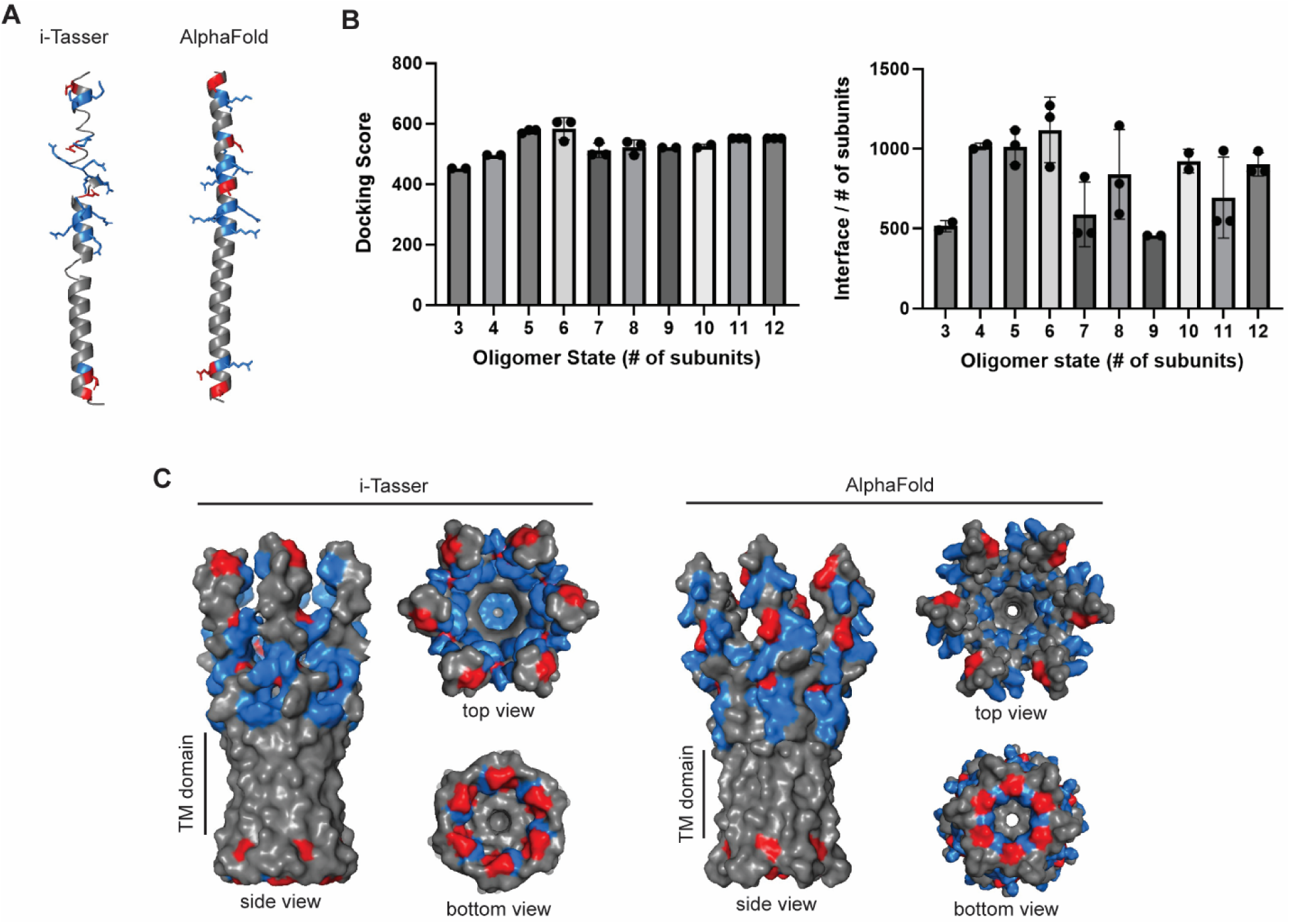
Simulation of Mtln self-association predicts stable hexamer formation. **A.** Singlet models of human Mtln generated using default settings on i-Tasser and AlphaFold show linear alpha-helical structures. Models were visualized using PyMOL software. Negatively- and positively-charged amino acids are indicated by red and blue respectively. **B.** Summarized average docking scores and normalized interface areas (interface/number of subunits) are plotted for each possible oligomer based on simulations from GALAXYHomomer. Individual datapoints for the three singlet models (Alphafold and two i-Tasser models)) analyzed are plotted. **C.** Structural models of human Mtln hexamers generated from GALAXYHomomer using singlet i-Tasser and AlphaFold models. Models were visualized using PyMOL. Transmembrane (TM) spanning portions are marked.

### Native-state analyses reveal Mtln complex formations consistent with a hexameric form

To reexamine the size of possible Mtln-containing multimeric complexes in wet-lab studies, we isolated mitochondria from fresh or snap-frozen wildtype (WT) and Mtln-KO A549 cells and performed BN-PAGE, given that Zhang et al.^12^ found that freezing induces Mtln aggregation (**Fig. 3A**). As previously observed in mouse heart and skeletal muscle mitochondrial isolates^5,12^, a prominent Mtln-containing complex migrates at ∼66 kDa in WT samples, and this band is absent from KO samples and decreased in frozen samples with coinciding accumulation of higher molecular weight aggregates. Of note, this 66 kDa complex does not appear to be merely migrating at the protein/dye front in the BN-PAGE, as is likely the case for the fainter band below, which we suspect corresponds to singlet Mtln.

**Figure 3.**
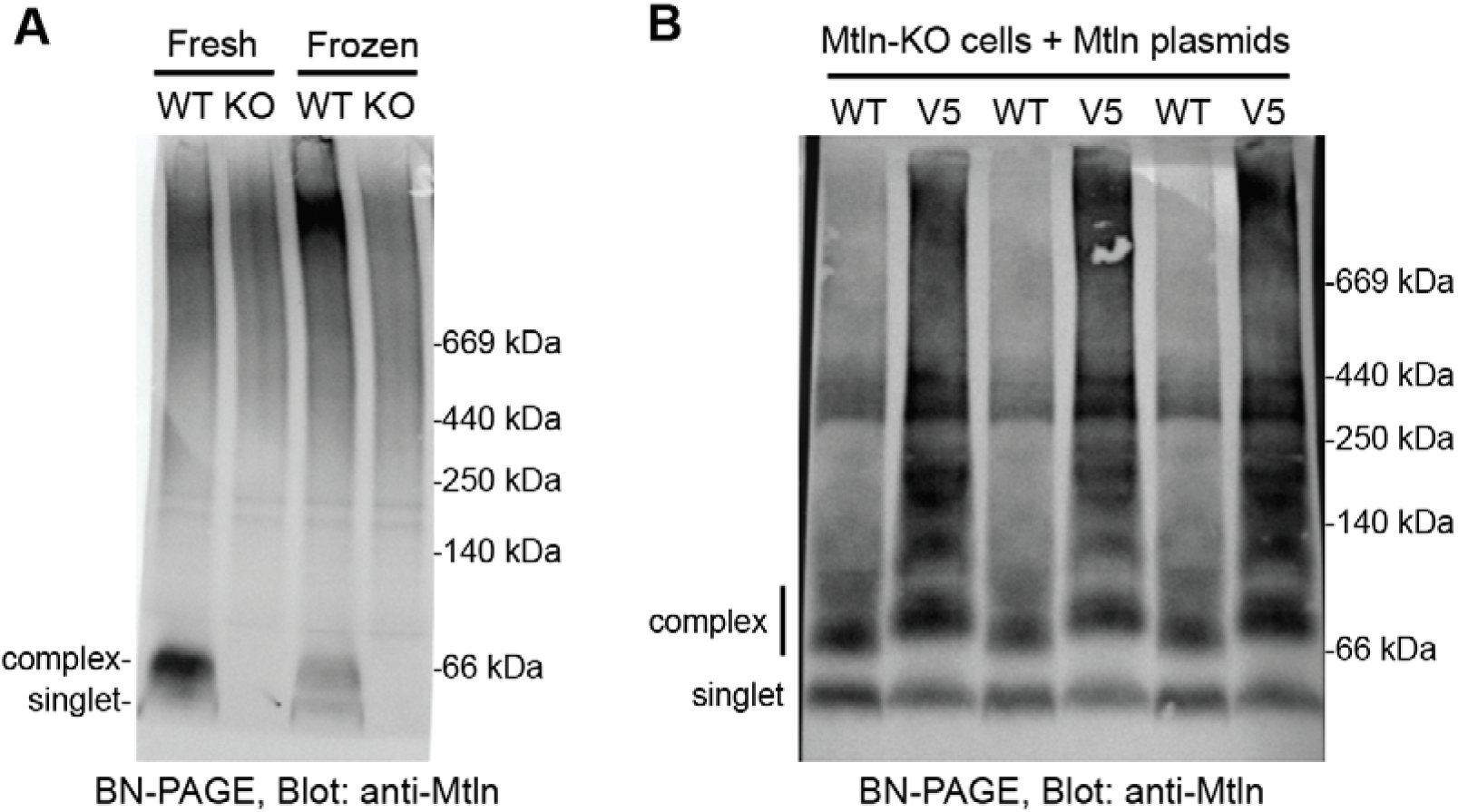
Endogenous and overexpressed Mtln is present in a prominent 66 kDa complex consistent with hexameric Mtln. **A.** BN-PAGE (3-12% gel) assessment of endogenous Mtln in mitochondria collected from fresh or frozen wildtype (WT) or Mtln-KO A549 cells. **B.** Native Mtln or V5-tagged Mtln were expressed in Mtln-KO A549 cells, and their abilities to form / integrate into complexes were assessed by BN-PAGE (4-16%). Bands corresponding to presumed Mtln singlets and hexamer complexes are denoted.

Given our observations of Mtln self-association in co-immunoprecipitation experiments and computational simulation data supporting a possible Mtln hexameric structure, we conducted additional studies to gain insight into whether the band at 66 kDa might be a homo-oligomer of six Mtln subunits, each of which is ∼11 kDa alone (based on SDS-PAGE migration). We transfected Mtln-KO A549 cells with WT or V5-tagged Mtln expression plasmids and conducted BN-PAGE on fresh mitochondrial isolates to assess if and to what extent V5-tagged Mtln complexes shift to higher molecular weights relative to WT Mtln complexes (**Fig. 3B**). Notably, V5-tagged complexes shift up to ∼75 kDa, representing a 9 kDa increase in complex size relative to WT Mtln complexes, which could be accounted for by the addition of six 1.5 kDa V5 tags, consistent with formation of a hexameric Mtln complex.

### Mtln aggregates to form higher-order oligomeric SDS-resistant complexes in fresh tissues

In previous BN-PAGE analyses done on mouse skeletal muscle mitochondria isolates, Zhang et al. showed a predominant ∼66 kDa band and low levels of higher molecular weight smearing of Mtln signal in fresh samples, which was almost completely inversed in intensities in frozen samples, prompting the conclusion that freezing artifact promotes Mtln aggregation^12^. These findings are consistent with our observations in A549 cells (**Fig. 3A**). To further explore the prospect of Mtln to form multimeric or aggregated forms *in vivo,* we examined Mtln-containing complexes in mouse cardiac tissue samples, as *in silico* structural modeling also predicted that mouse Mtln could form strongly docked and highly interfaced hexameric complexes (**Fig. S4**). We performed BN-PAGE on mitochondrial isolates collected from fresh and snap-frozen WT or Mtln-KO mouse hearts (**Fig. 4A**). Additionally, 1^st^ dimension BN-PAGE was followed by 2^nd^ dimension SDS-PAGE to resolve Mtln to its singlet/monomeric form and identify SDS-insoluble forms that fail to appreciably enter the second gel (**Fig. 4B**). In 1^st^ dimension BN-PAGE blots, we found that frozen samples clearly showed less Mtln signal at ∼66 kDa and more robust accumulation of putative Mtln aggregates smearing at higher molecular weights. In fresh samples, we surprisingly observed quite strong Mtln signal in high molecular weight smears, as well as distinct lower intensity bands at ∼66 kDa and below, likely corresponding to oligomeric and singlet Mtln respectively. We considered the possibility that the lower signal intensity at ∼66 kDa could be due to limited antibody epitope accessibility in these native 66 kDa Mtln complexes. Indeed, when resolved in 2^nd^ dimension SDS-PAGE (**Fig. 4B**), the majority of Mtln (11 kDa in the 2^nd^ dimension, spot marked by red arrow) aligns with a ∼66 kDa point of origin from the 1^st^ dimension. While it is likely that singlet Mtln (black arrow) also contributes to signal intensity of this spot, the center of the spot appears to align best with the 66 kDa complex (red arrow). Of note, this 2^nd^ dimension Mtln-positive spot was absent from KO samples and frozen WT samples, providing further evidence that freezing disrupts the 66 kDa complex and promotes Mtln aggregation. Nevertheless, it is important to note that, even in fresh samples, a considerable amount of singlet Mtln resolved from 1^st^ dimension high molecular weight smears (solid box outline), suggesting that these higher-order complexes are not merely the result of freezing artifact. Furthermore, we noted the presence of SDS-resistant, Mtln-containing higher molecular weight complexes of various sizes, evidenced by apparent laddering from ∼120 to 260 kDa in the 2^nd^ dimension (dotted box outline). Although these complexes were more abundant in frozen samples, consistent with some freezing artifact, reasonable levels were also detected in fresh samples (most prominently at ∼140 kDa), supporting their potential natural existence in heart mitochondria. It is possible that these ∼120 to 140 kDa complexes correspond to predicted 11-mer and 12-mer Mtln oligomeric complexes, which presented with reasonable membrane spanning widths in our structural modeling investigations (**Fig. S3**).

**Figure 4:**
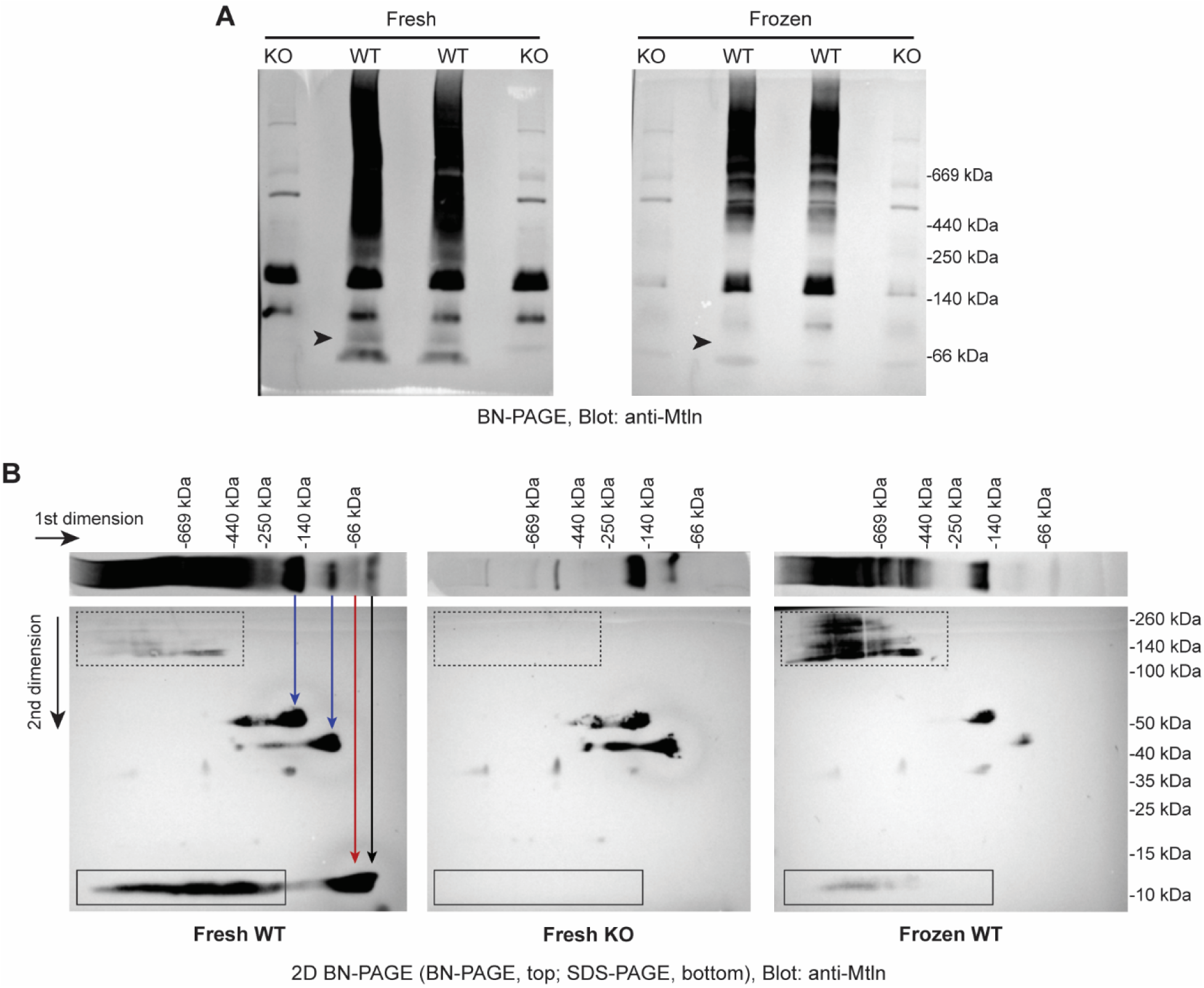
Mtln is present in a prominent 66 kDa complex and in higher-order complexes in fresh mouse heart tissues. **A.** BN-PAGE assessment of mitochondrial isolates collected from fresh or frozen wildtype (WT) or Mtln-KO mouse heart tissues. Arrowheads point to the region corresponding to the 66 kDa complexes that appears to contain the bulk of Mtln (see panel B, red arrow). **B.** 2D BN- and SDS-PAGE of mitochondrial isolates collected from fresh or frozen WT and Mtln-KO mouse heart tissue. Example lanes from panel A are placed at top for alignment purposes, along with arrows (blue denoting two non-specific bands that are present in KO samples, red corresponding to the 66 kDa complex and black indicating singlet Mtln). Solid boxes outline regions of singlet Mtln derived from high molecular weight 1^st^ dimension smears in WT samples; dotted box outlines areas encompassing SDS-resistant Mtln complexes in WT samples.

### Synthetic Mtln self-associates to form complexes consistent in size to hexameric structures

Thus far, we have demonstrated that Mtln self-associates (by co-immunoprecipitation) and predominantly resides within a 66 kDa complex (by BN-PAGE). While this size fits with a hexameric Mtln complex predicted by *in silico* modeling, it is possible that these are heteromeric protein complexes, and it remains unknown whether Mtln alone can self-oligomerize. To address this, we obtained synthetic Mtln protein and performed BN-PAGE analyses on solubilized protein with and without addition of detergent. While Mtln alone did not produce a clear high molecular weight band, an apparent 66kDa Mtln complex was formed in the presence of 1% digitonin, which provides a lipid-like environment and supports oligomeric structures (**Fig. 5A**). In complementary experiments, we performed dynamic light scattering (DLS) to determine the hydrodynamic radius distribution of solubilized Mtln. The measured size distribution by mass gave a single peak at 3.8 ± 0.06 nm which is calculated to be 76.5 ± 3.1 kDa (**Fig. 5B**). Although this is close to our observed 66 kDa complex on BN-PAGE, it is important to note that DLS can only give crude information regarding the size of molecules in solution, as the calculations are shape-dependent while assuming both a spherical protein structure and homogenous population of complexes^18^. To circumvent this, we also analyzed synthetic Mtln by mass photometry (MP), a method which is shape-independent and can provide sizes of different species of complexes in a mixture^19^. MP revealed a predominant peak at ∼70kDa, consistent with a potential Mtln hexamer, with slight “smearing” towards higher molecular weight complexes (**Fig. 5C**). Monomeric Mtln could not be measured by MP because it is below the range of detection (cannot typically quantify proteins less than ∼40kDa). Overall, these combined data support that synthetic Mtln is capable of forming homo-oligomeric complexes, with the predominant species matching the ∼66 kDa size of native Mtln-containing complexes found in mitochondrial isolates collected from cells and mouse tissues.

**Figure 5:**
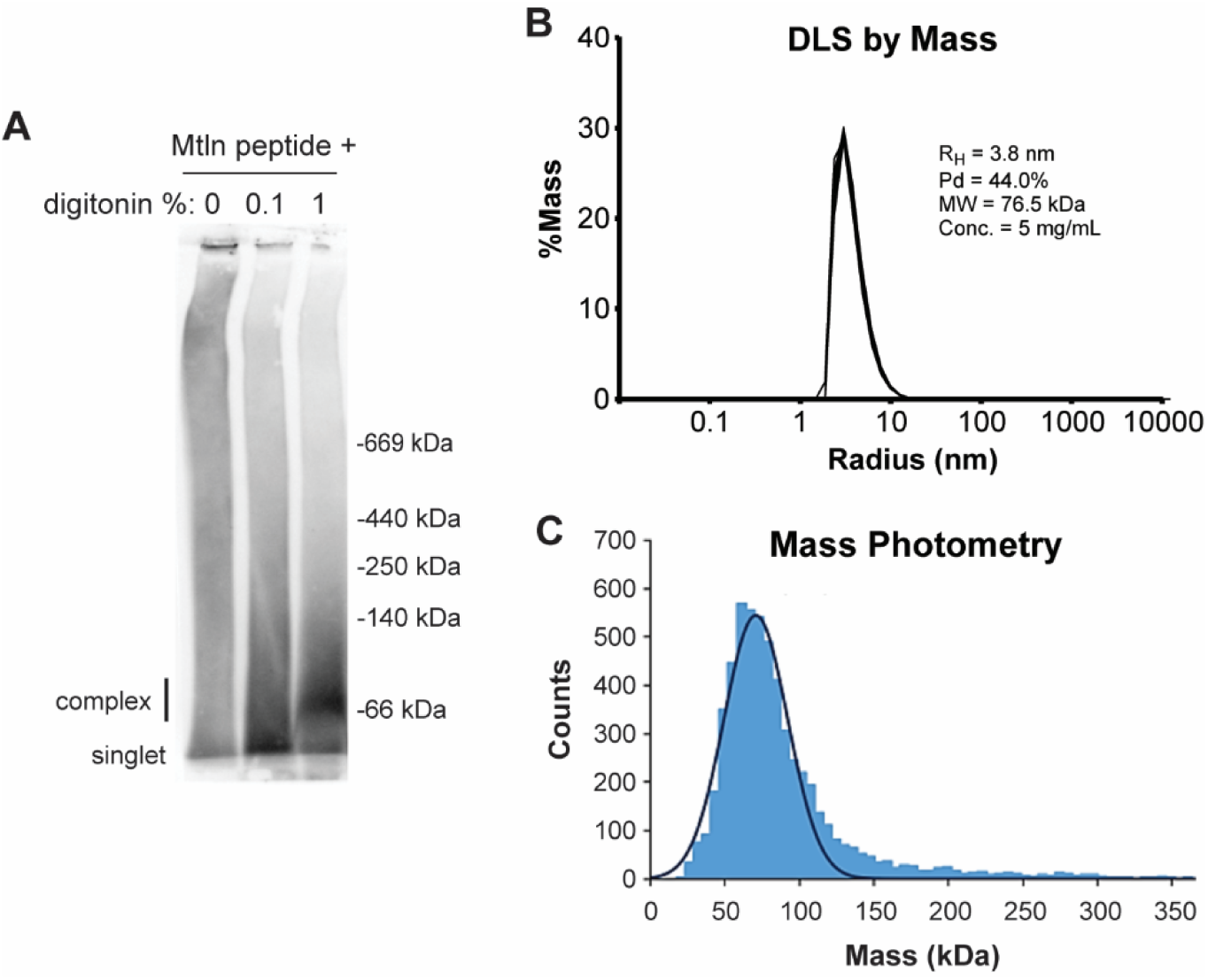
Synthetic Mtln protein self-associates to form complexes consistent in size to hexameric structures. **A.** Synthetic Mtln protein was solubilized with varying amounts of digitonin and visualized by BN-PAGE, with western blotting using anti-Mtln antibody. **B.** Dynamic light scattering (DLS) performed on synthetic Mtln in solution suggests the presence of a 76.5 kDa oligomeric complex by mass analysis. **C.** Mass photometry (MP) performed on solubilized synthetic Mtln indicates a prominent oligomeric complex of ∼70 kDa, with some possible “smearing” towards higher molecular weights.

## DISCUSSION

To date, many studies have examined the diverse functionality of Mtln, but no investigations have explored the possibility of Mtln homo-oligomerization. Through a comprehensive investigation of the possible native state of Mtln, we found that Mtln interacts with itself and forms higher-order complexes, including a predominant complex that is likely composed of six Mtln subunits. Our combined approaches leveraging co-immunoprecipitation, BN-PAGE done on human cell line and mouse tissue samples, analysis of synthetic Mtln by gel electrophoresis, DLS, and MP, along with simulated modeling all support the general conclusion that Mtln self-associates to form homo-oligomeric structures, prompting future studies to further characterize the precise nature and molecular actions of these complexes.

Mtln self-association was particularly evident (though perhaps overlooked) in co-immunoprecipitation studies done previously by Xiao et al.^9^, which we confirmed in our own experiments using a different cell line and a different epitope tag. Our studies in Mtln-KO cells were unique, removing the confound that endogenous (WT) Mtln may be pre-complexed and less likely to engage with transgene-derived tagged Mtln. By simultaneously co-expressing WT and V5-tagged Mtln and subsequently performing anti-V5 co-immunoprecipitation, we found that WT and nV5-Mtln were pulled down at nearly a 1:1 ratio, which supports a striking degree of co-interaction. This can also be inferred from the Xiao et al. data that clearly indicates that endogenous WT Mtln was co-immunoprecipitated with Mtln-FLAG at ∼5-10 times greater efficiency compared to their reported co-interactor, Ndufa7 (comparing blot data for their input and pulldown samples). It is possible that most prior investigations of Mtln interactors by co-immunoprecipitation coupled with mass spectrometry have been confounded by their reliance on supraphysiologic overexpression of epitope-tagged Mtln constructs delivered to cells that already express native Mtln. This could lead to unnatural forced interactions and pulldown of non-interacting mitochondrial membrane proteins within protein-detergent micelles. Indeed, many of the proposed Mtln-interacting proteins (from pulldown studies of both tagged and untagged Mtln) are known to be among the most highly abundant mitochondrial proteins. Nevertheless, our goal is not to negate the possibility for alternative (and likely weaker) interactions with other proposed binding partners such as Hadha/b, Cyb5r3, Ndufa7 and Cpt1b, but rather to present data supporting that Mtln is primarily a self-interacting protein that can homo-oligomerize into complexes with unknown molecular function, with overall hopes to inspire focused studies to further explore this.

In characterizing Mtln self-association, our study identifies a collection of six subunits (hexamer) as the most likely homomeric structure. Hexameric formation is also supported by our simulated modeling investigations, indicated by the highest docking score and normalized interface area that we observed across the >30 possible models generated. An important aspect and potential limitation of utilizing GALAXYhomomer is its lack of consideration for transmembrane protein constraints. Thus, we only considered output models that exhibited parallel and aligned transmembrane domains that could plausibly span mitochondrial membrane widths (∼3 nm). Beyond Mtln hexamer models, which we focused on given the overall conglomerate of our observations in wet-lab studies, several other models with varying numbers of subunits could plausibly span membranes. We certainly cannot rule out the possible natural existence of these complexes, and notably, Mtln-positive bands migrating at 120-260 kDa could correspond to dodecameric and/or other higher-order Mtln complexes, some of which were also predicted to be strongly docked and highly interfaced with adequate transmembrane spanning distances.

It is possible that Mtln exists in various concurrent states or alternates between states. In our BN-PAGE plus 2^nd^ dimension SDS-PAGE assessments of mouse heart mitochondrial isolates, we noted that singlet Mtln arises not only from the 66 kDa complex, but also migrates out of complexes spanning a range of higher molecular weights. This differs from skeletal muscle^12^ or human cell lines (**Fig. 3A**), where fresh isolates primarily generate the 66 kDa form. It is unclear whether the discrepant findings could arise from subtle changes in sample processing and BN-PAGE protocols or represent tissue-specific variations in mitochondrial compositions and Mtln complexes. For the latter, we consider a scenario in which Mtln may integrate into and/or associate with higher-order complexes within the protein-rich and cardiolipin-rich membrane microdomains in cardiomyocyte mitochondria.

At this time, we can only speculate on possible functions of the Mtln homo-oligomeric complexes. Given its transmembrane domain and possible localization to both inner and outer mitochondrial membranes, Mtln may hexamerize to form a pore, channel, or transporter of some capacity, which could provide an avenue through which Mtln influences membrane potential, ROS and/or calcium handling. Indeed, there are several published instances of mitochondrial proteins that homo-oligomerize to function in these facets, as well as in fostering cristae formation / stability^20–23^. Given Mtln’s known tendency to strongly bind cardiolipin, Mtln oligomers could act as structural scaffolds to arrange mitochondrial membranes and/or protect cardiolipin to support an array of mitochondrial processes, including respiration via the electron transport chain. Future investigations could focus on identifying Mtln mutations or peptides that block homo-oligomerization to begin teasing out the molecular action and physiologic relevance of these complexes or whether some Mtln functions can be attributed to singlet forms. Along these lines, we speculate the Mtln self-association might depend upon a tryptophan residue (Trp25) in the transmembrane domain, given a breadth of literature supporting key roles for tryptophans in mediating protein-protein interactions, particularly in transmembrane helices^24,25^. In addition, studies should address whether Mtln oligomerization is disrupted in and/or influences disease (e.g. mitochondrial disorders, neurodegenerative diseases, or cancer), with some consideration for possible therapeutic implications of developing small molecules that modulate Mtln complex formation and function.

In summary, we establish that Mtln self-associates and likely forms a stable homo-oligomeric hexamer. Follow-up studies will be needed to definitively validate that Mtln, as we suspect, binds to itself with much higher affinity compared to its many other proposed protein interactors. Beyond human cell lines and mouse muscle tissues, it will be important to examine the presence of Mtln homo-oligomeric complexes across other tissues and species, including fresh human muscle tissues. Finally, while we present a breadth of initial complementary observations that support Mtln homo-oligomerization, more advanced studies will be needed to empirically define the resulting complexes at the structural level. Mtln hexameric complexes are potentially too small to be defined by cryo-EM, but perhaps further insights could be gained by performing negative stain EM on Mtln complexes reconstituted in lipid bilayers (e.g. nanodiscs).

## LIMITATIONS OF THE STUDY

A limitation of this study is that our *in silico* protein structure predictions were not done in the context of a modeled membrane, and thus, do not account for possible high affinity interactions between Mtln and cardiolipin, which could likely influence the nature of Mtln self-association. Additionally, although we confirmed tagged-Mtln localization to mitochondria, we cannot confirm whether the presence of the V5 epitope impacts Mtln orientation and function, though Mtln is thought to exist in both inner and outer membranes. Finally, our findings (along with previously unrecognized data form Xiao et al.^9^) support that Mtln’s most prominent protein interactor is likely itself; however, our proposed Mtln homo-oligomeric complexes and structures remain to be fully resolved.

## ACKNOWLEDGEMENTS

This work was supported by the University of Iowa Carver College of Medicine (Distinguished Scholars Program to R.L.B.), NIH NHLBI (HL150557 to R.L.B.), NIH NIGMS (predoctoral fellowship T32 GM067795 to N.H.W.), American Heart Association (20IPA35360150 to R.L.B. and 23PRE1011277 to N.H.W.), and University of Iowa Office of Undergraduate Research (fellowship to C.R.L.). We also acknowledge the University of Iowa core facilities that made significant contributions to this work, including the Protein and Crystallography Facility, Genomics Division, and Microscopy Core.

## AUTHOR CONTRIBUTIONS

R.L.B. conceived the project, supervised research, and analyzed and interpreted data. C.R.L., C.S.S., and N.H.W. designed and executed experiments and analyzed and interpreted data. Z.X. and N.J.S. executed experiments. R.L.B., C.R.L., and N.H.W. wrote the manuscript with contributions from Z.X. and N.J.S.

## SUPPLEMENTAL INFORMATION

Document S1. Figures S1–S4.

## MATERIALS AND METHODS

### Cultured Cell Lines

Human A549 (ATCC, CCL-185) cells were maintained in DMEM/F12 with 100U/ml penicillin, 0.1mg/ml streptomycin, and 10% fetal bovine serum, in a cell culture incubator under standard conditions (37°C in 5% CO_2_). Cells were kept at a low passage and regularly passaged prior to 100% confluency.

### Generation of CRISPR/Cas9 Mtln knockout A549 cell line

CRISPR sgRNA expression plasmids were generated to target regions upstream and downstream of the human Mtln open reading frame. Five upstream and five downstream guide sequences were cloned into an expression vector (pFBAAV-mU6-sgRNA-CMV-eGFP-SV40pA). 25 up-down guide pairs were tested for cutting efficiency in A549 cells by co-transfection with an spCas9 expression plasmid (pU6-BbsI-CBh-Cas9-T2A-mCherry; Addgene #64324). Cutting efficiency was evaluated after 48hrs by performing PCR on gDNA using primers flaking the Mtln ORF (data not shown). For cell line generation, the two most efficient guide pairs were co-transfected with Cas9 and after 48hrs, the cells were lifted and sorted to enrich for dually-transfected cells and seeded at one cell per well in a 96-well plate. The resulting clones were screened by PCR (as described above) for Mtln gDNA knockout. Mtln knockout was confirmed at the protein level by western blot. Primers and sgRNA sequences are listed in Table 1.

**Table 1:**
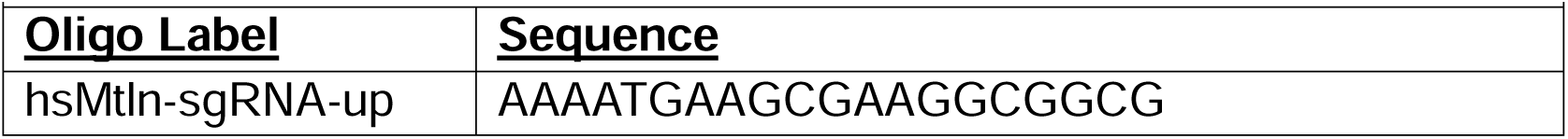
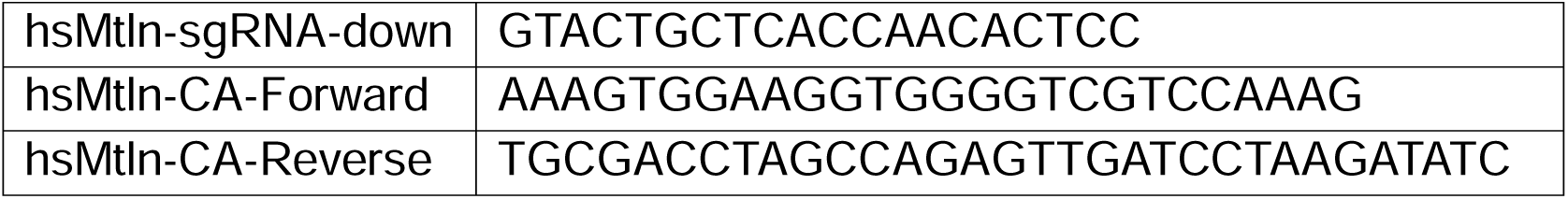
DNA oligo sequences.

### Plasmids and Transfection

A549 wildtype or knockout cells were transfected with various human Mtln expressing plasmids (wildtype, N-terminal or internal V5 epitope-tagged) using Lipofectamine LTX (Thermo Fisher Scientific) according to the manufacturer’s instructions. 48-72 hours post-transfection, cells were assayed. Detailed plasmid maps are available upon request to the corresponding author.

### Cell Staining

Mtln-KO A549 cells were seeded on glass-like plates and transfected with the indicated plasmids using Lipofectamine LTX according to the manufacturer’s instructions. After 48 hours, live cells were incubated with MitoTracker™ Deep Red and then washed 3x with PBS. Cells were fixed with 4% PFA and blocked/permeabilized in a PBS solution with 0.1% Triton and 5% goat serum. Cells were then incubated with custom anti-Mtln primary antibody overnight (1/500). Cells were washed and incubated with indicated secondary antibody (Alexa Flour). Cells were washed and visualized by confocal microscopy, and images were analyzed using ImageJ.

### Immunoprecipitation

60mm-dishes of transfected cells were harvested in lysis buffer (12mM NaCl, 50mM Tris, 1mM EDTA, 1% NP-40, pH 7.5) containing protease/phosphatase inhibitors and clarified by centrifugation. 50ul of suspended V5-Trap Magnetic Agarose (ChromoTek) was prepared according to the manufacturer’s instructions and added to each lysate and incubated for 2 hours at 4°C with gentle continuous inversion. The agarose was washed 4 times, 10 minutes each using lysis buffer containing 0.1% NP-40. Proteins were eluted in sample buffer (NuPAGE LDS and Reducing Agent) at 80°C for 10 minutes. Input and depleted fractions were also collected.

### Western Blotting

Lysates were prepared in NuPAGE LDS and Reducing Agent with heating at 80°C for 10 minutes and resolved on 4-12% Bis-Tris gels using NuPAGE MES running buffer. Proteins were transferred to 0.2um PVDF membranes using NuPAGE transfer buffer with 10% ethanol. The membrane was blocked with 5% milk in TBST (0.1% Tween-20) and incubated in the indicated primary antibody overnight at 4°C. Membranes were washed with TBST, incubated for 1 hour at room temperature with HRP-conjugated secondary antibody, washed, and visualized using Femtogram HRP substrate (Azure Biosciences) on an iBright 1500 (Thermo Fisher Scientific).

### Mitochondrial Isolation

#### Cultured cells

Method adapted from Frezza et al.^26^. 10cm dishes of cells were washed, scraped into PBS, and pelleted. For frozen samples, cells were washed and dishes floated in liquid nitrogen prior to scraping and pelleting cells in PBS. Fresh or frozen cells were then resuspended in 300ul mitochondrial isolation buffer (200mM sucrose, 10mM Tris, 1mM EGTA, pH 7.4) and homogenized in a 2ml dounce tissue grinder (Kimble) with 40 passes using “pestle B”. Intact cells and debris were removed by centrifugation at 800xg for 5 minutes. The supernatant was then centrifuged at 8,000xg for 10 minutes to pellet mitochondria.

#### Mouse heart

The heart was excised from wildtype or Mtln knockout (Mtln-KO, previously described^5^ mice immediately after euthanasia and rinsed in ice-cold PBS to remove excess blood. For frozen samples, half of the heart was snap frozen in liquid nitrogen and thawed on wet ice prior to mitochondria isolation. Heart tissue was minced into <1mm^2^ pieces and washed in STE-1 buffer before homogenization in a size 22 tissue grinder (Kontes) in 4.5mL STE-2 buffer. Tissue debris were removed by centrifugation at 800xg for 10 minutes, and mitochondria were pelleted from the resulting supernatant by centrifugation at 8,000xg for 10 minutes.

### Blue Native PAGE

Blue Native PAGE procedures were derived from previously described methods^5^. Briefly, mitochondria isolates were immediately resuspended in ACA buffer and proteins quantified (BCA assay, Thermo Fisher Scientific) and equalized between samples. Mitochondrial proteins were solubilized with 1% digitonin resolved on 4-16% or 3-12% NativePAGE Novex gels (Invitrogen). For <u>blotting</u>, proteins were transferred to 0.2um PVDF membranes, fixed with 8% acetic acid for 10 minutes and dried overnight. Membranes were washed with methanol to remove the blue dye and equilibrated with water before proceeding with standard western blot procedures. For <u>second dimension analysis</u>, lanes were excised from the NativePAGE gels, trimmed, and denatured in NuPAGE LDS sample buffer containing reducing agent for 30 minutes at room temperature. Lanes were fit horizontally onto NuPAGE 4-12% Bis-Tris gels with a 2D well, overlayed with sample buffer, and resolved using NuPAGE MES running buffer. Proteins were transferred to membranes and treated as a standard western blot. For <u>synthetic Mtln</u> analysis, lyophilized protein was dissolved in water and diluted to final 10ug/ml in buffer (250mM sucrose, 10mM HEPES, pH 7.5). The indicated amount of digitonin was added, and incubated for 30 minutes at room temperature before proceeding with Blue Native PAGE blotting.

### Structure Simulations

Singlet models of Mtln were generated using default settings on iTasser (https://zhanggroup.org/I-TASSER/) and AlphaFold (https://alphafold.ebi.ac.uk/). The top two models from iTasser were chosen, along with the top model from AlphaFold. Each model was entered into GALAXYhomomer and models from 3-12 subunits were analyzed. Homomers of only 2 subunits were omitted due to their anti-parallel orientation. Normalized interface score was calculated by dividing interface score by the number of subunits.

### Dynamic Light Scattering (DLS)

Synthetic Mtln was dissolved in a buffer (50 mM NaAc, pH 4.0, 50 mM NaCl) with a concentration of 5 mg/mL. Sample solution was centrifuged at 18,300g for 25 minutes before measurements. The microcuvette (Wyatt Technologies, P/N: WNDMC) was filled with 12 μL Mtln solution. The DLS measurement were performed using DynaPro NanoStarTM (Wyatt Technologies) at 25 °C. All DLS data were analyzed by the DYNAMICS software (Wyatt Technologies) to calculate the hydrodynamic radius (RH) distribution.

### Mass Photometry (MP)

MP experiments were performed on a Refeyn TwoMP mass photometer (Refeyn Ltd, Oxford, UK). Microscope coverslips (24 mm x 50 mm, Thorlabs Inc.) were cleaned by serial rinsing with Milli-Q water and HPLC-grade isopropanol (Sigma Aldrich) followed by drying with a filtered air stream. Silicon gaskets (Grace Bio-Labs) to hold the sample drops were cleaned in the same procedure immediately prior to measurement. All MP measurements were performed at room temperature using Dulbecco’s phosphate-buffered saline (DPBS) without calcium and magnesium (Thermo Fisher). The instrument was calibrated using a protein standard mixture: β-amylase (Sigma-Aldrich, 56, 112 and 224 kDa), and thyroglobulin (Sigma-Aldrich, 670 kDa). Before each measurement, 15 µL of DPBS buffer was placed in the well to find focus. The focus position was searched and locked using the default droplet-dilution autofocus function after which 5 µL of Mtln at 6 uM was added and pipetted up and down to briefly mix before movie acquisition was promptly started. Movies were acquired for 60 s (3000 frames) using AcquireMP (Refeyn Ltd) using standard settings. All movies were processed, analyzed using DiscoverMP (Refeyn Ltd).

---

**Figure S1:**
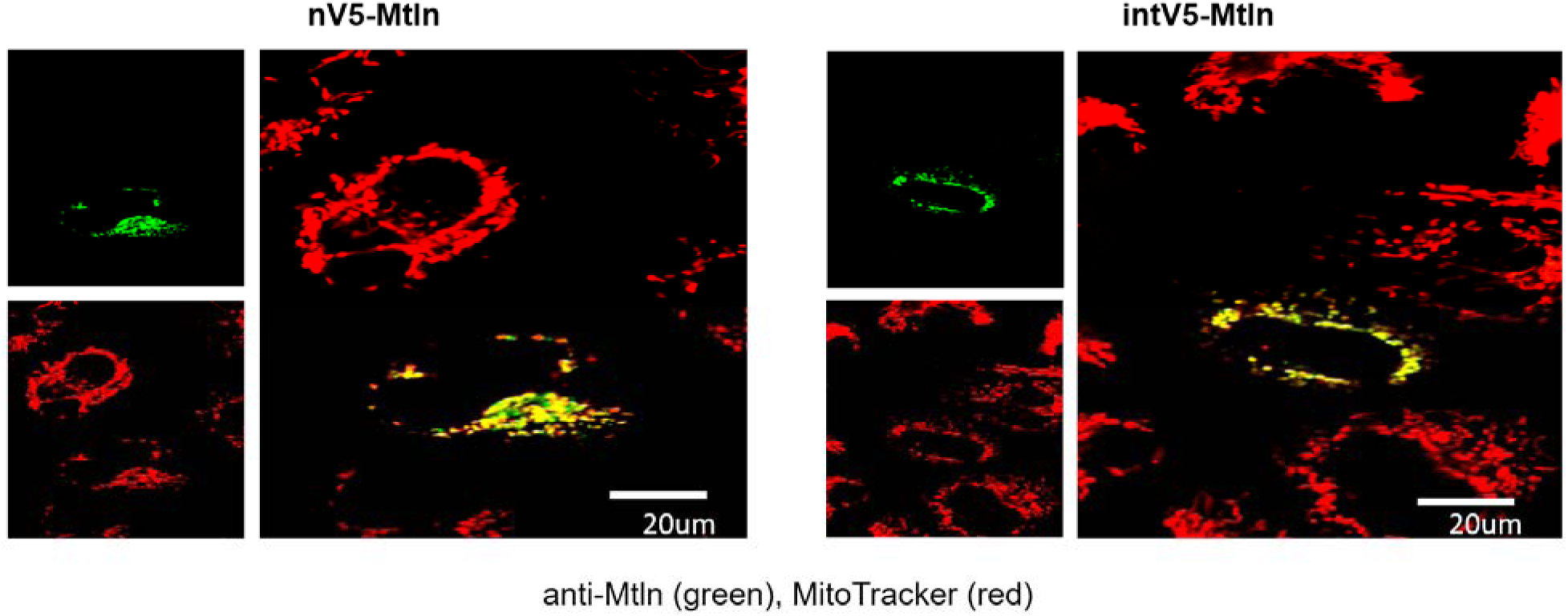
V5-tagged Mtln proteins localize to mitochondria. V5-tagged Mtln expression plasmids were transfected into Mtln-KO cells. Mitochondria were subsequently labeled by MitoTracker™ (shown in red) and anti-Mtln immunostaining (shown in green), the latter of which is clearly absent in non-transfected KO cells. Merged overlay images are also shown for each construct.

**Figure S2:**
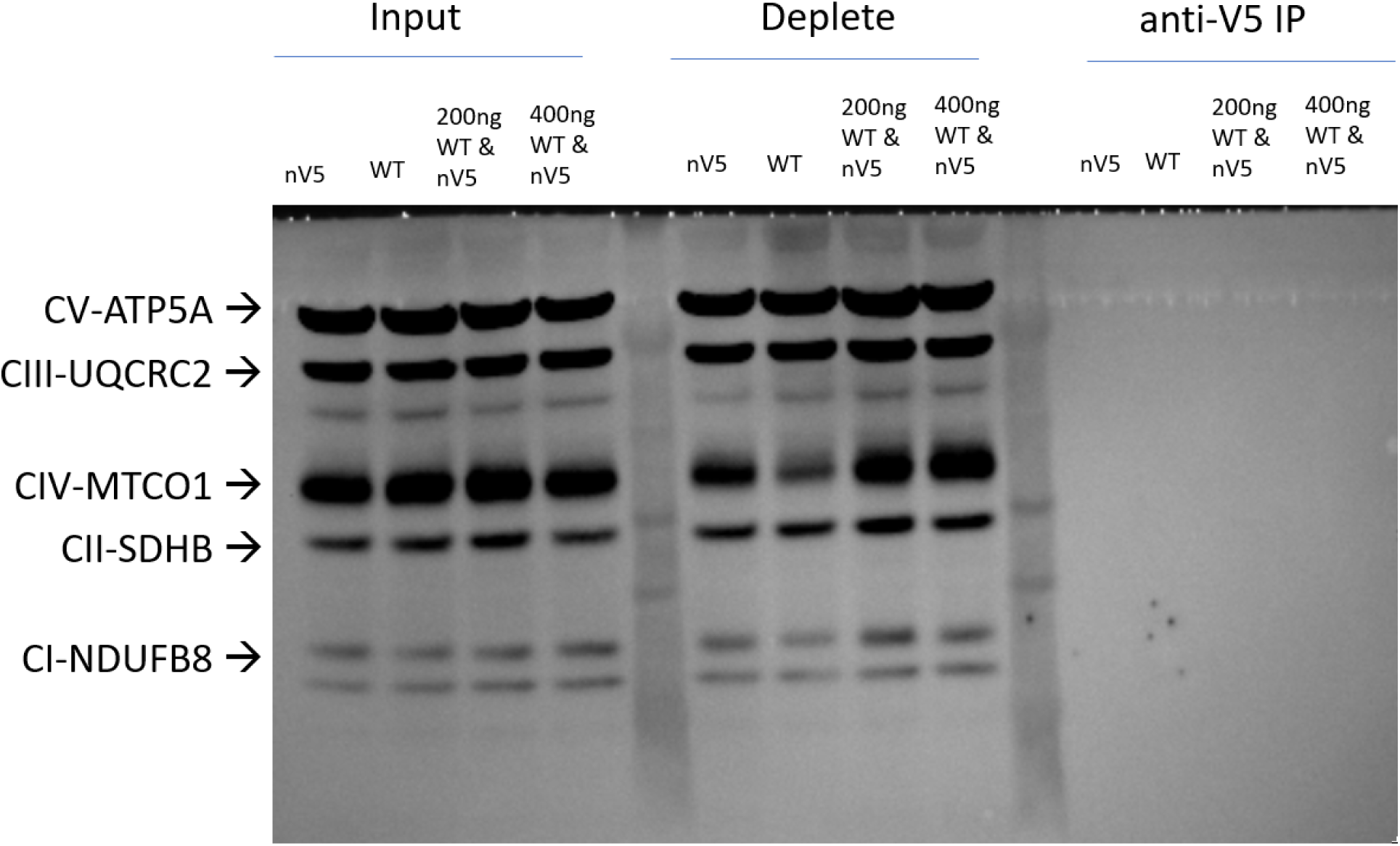
Anti-V5 immunoprecipitation does not generically pulldown mitochondrial membrane proteins. Input, deplete, and immunoprecipitation (IP) samples described in Figure 1D were blotted using an OXPHOS cocktail antibody.

**Figure S3.**
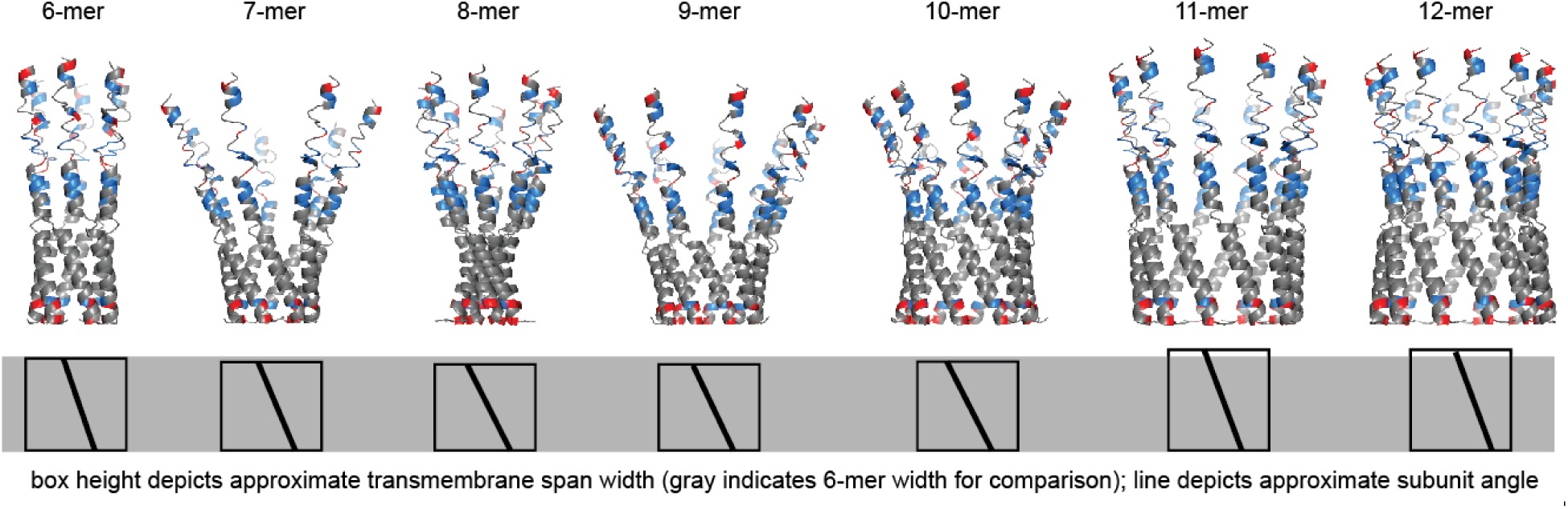
Hexameric (6-mer) and potentially 11- and 12-mer Mtln complexes have highly interfaced subunits and membrane domain lengths that could more plausibly span mitochondrial membrane widths. Possible human Mtln oligomeric complexes generated from i-Tasser singlet model input into GALAXYHomomer are shown with schematic representations depicting subunit angles and transmembrane domain spanning lengths (box heights). Gray shaded box indicates about ∼2.8 nm corresponding to the estimated hexameric (6-mer) transmembrane spanning width. Mitochondrial membrane widths are estimated to be ∼3 nm.

**Figure S4:**
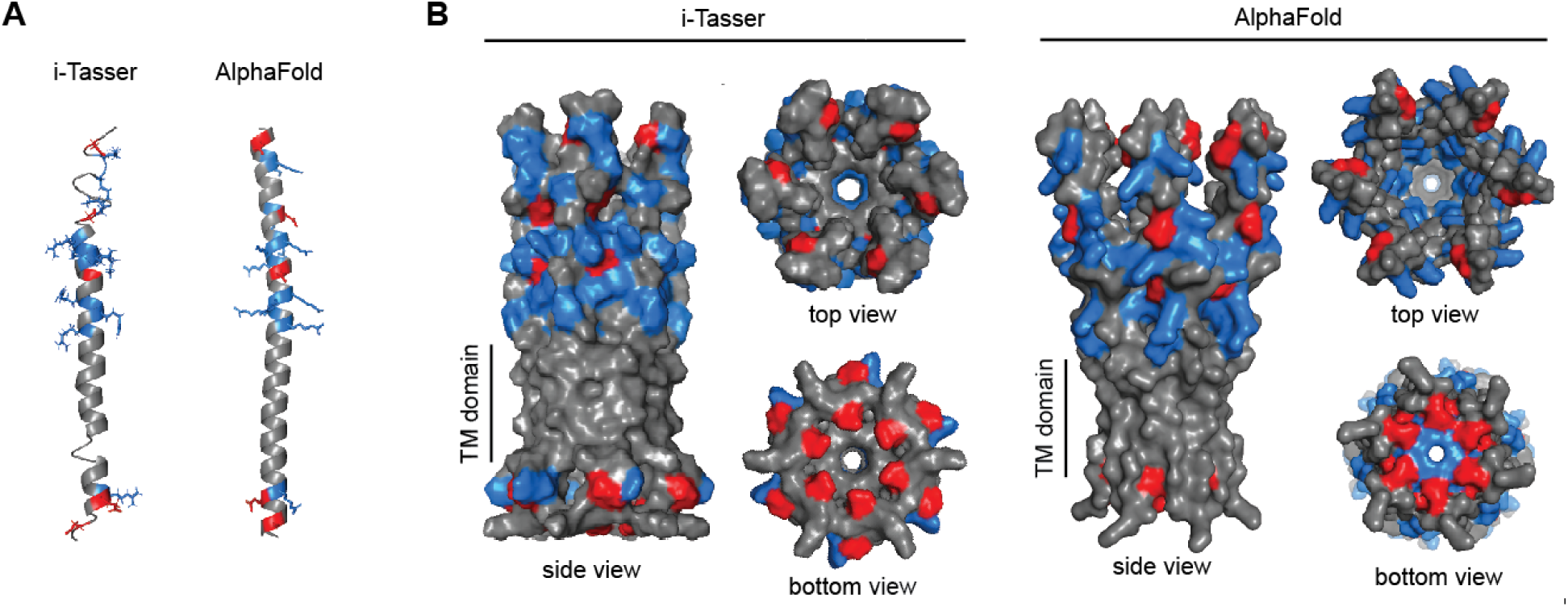
Mouse Mtln homo-oligomerization may form a hexameric pore-like complex. **A.** Singlet models of mouse Mtln generated using default settings on i-Tasser and AlphaFold show linear alpha-helical structures. Models were visualized using PyMOL software. Negatively- and positively-charged amino acids are indicated by red and blue respectively. **B.** Structural models of a mouse Mtln hexamers generated from GALAXYHomomer using singlet i-Tasser and AlphaFold models. Models were visualized using PyMOL. Transmembrane (TM) spanning portions are marked.

