## Supplemental Figures for "Mitoregulin self-associates to form likely homo-oligomeric pore-like structures"

**SUPPLEMENTAL INFORMATION**

**Figure S1**

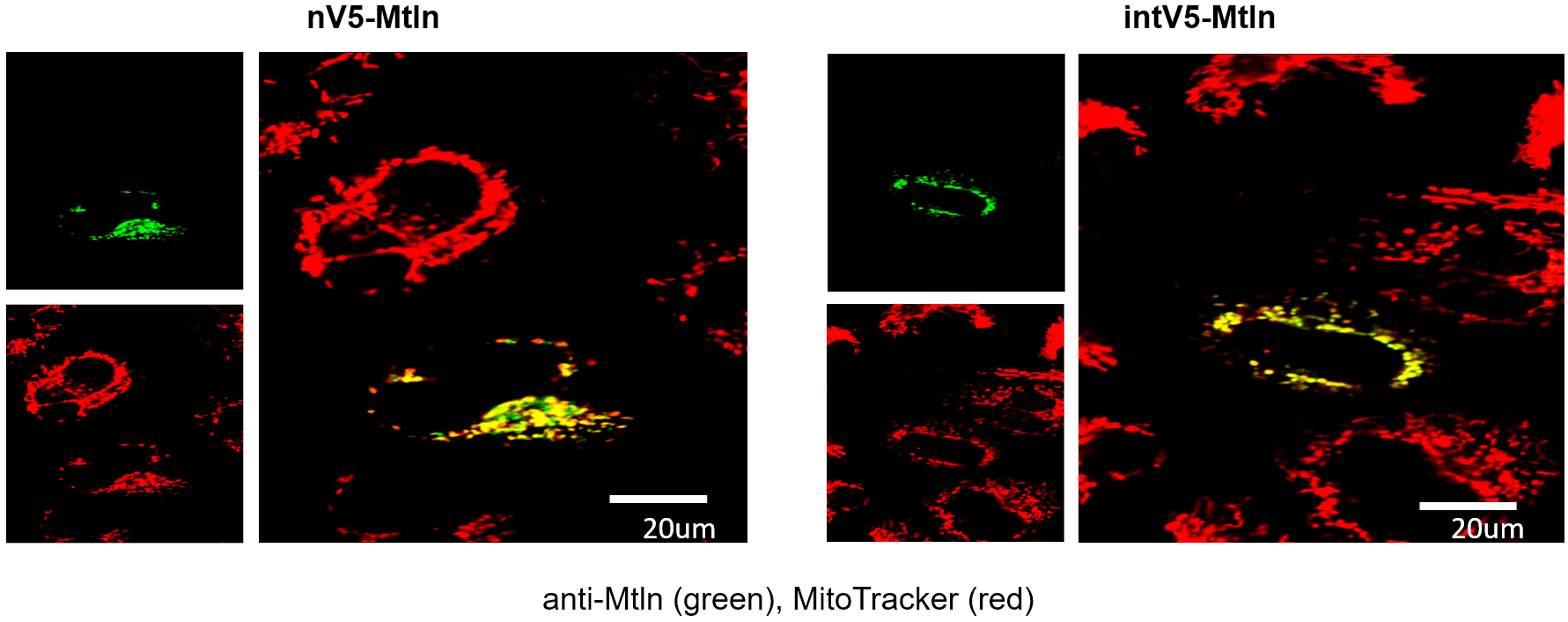

**Figure S1: V5-tagged Mtln proteins localize to mitochondria.**

V5-tagged Mtln expression plasmids were transfected into Mtln-KO cells. Mitochondria were subsequently labeled by MitoTracker™ (shown in red) and anti-Mtln immunostaining (shown in green), the latter of which is clearly absent in non-transfected KO cells. Merged overlay images are also shown for each construct.

**Figure S2**

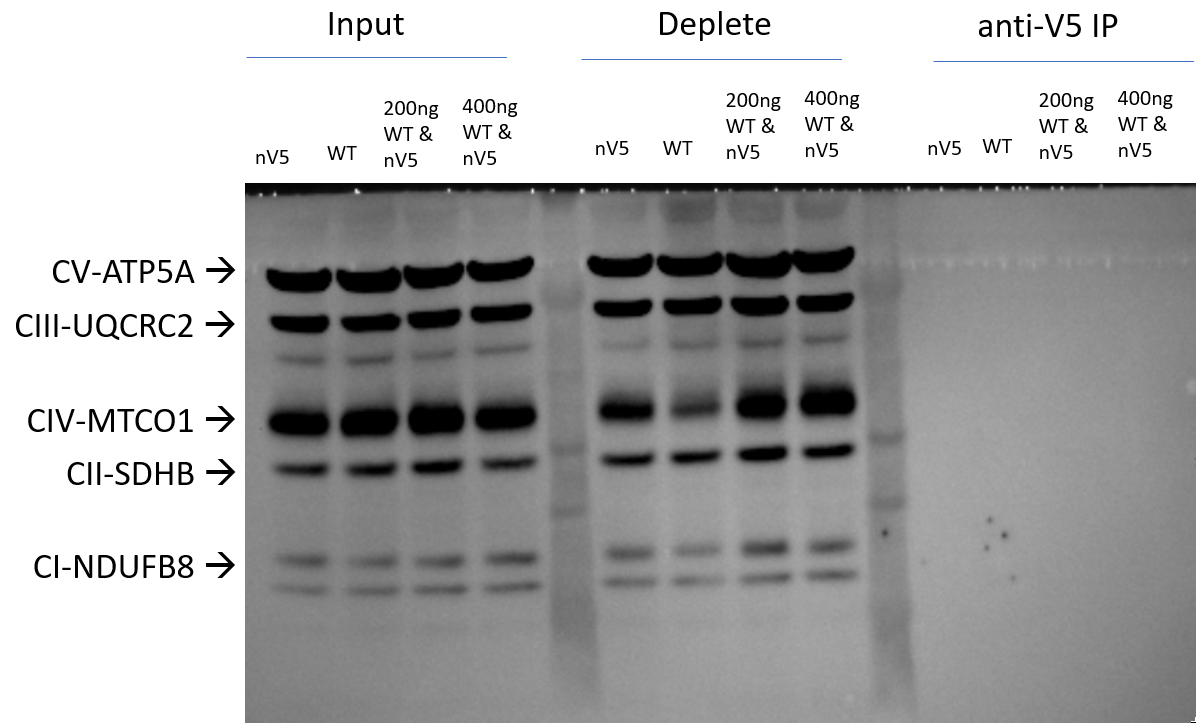

**Figure S2: Anti-V5 immunoprecipitation does not generically pulldown mitochondrial membrane proteins.**

Input, deplete, and immunoprecipitation (IP) samples described in Figure 1D were blotted using an OXPHOS cocktail antibody.

**Figure S3**

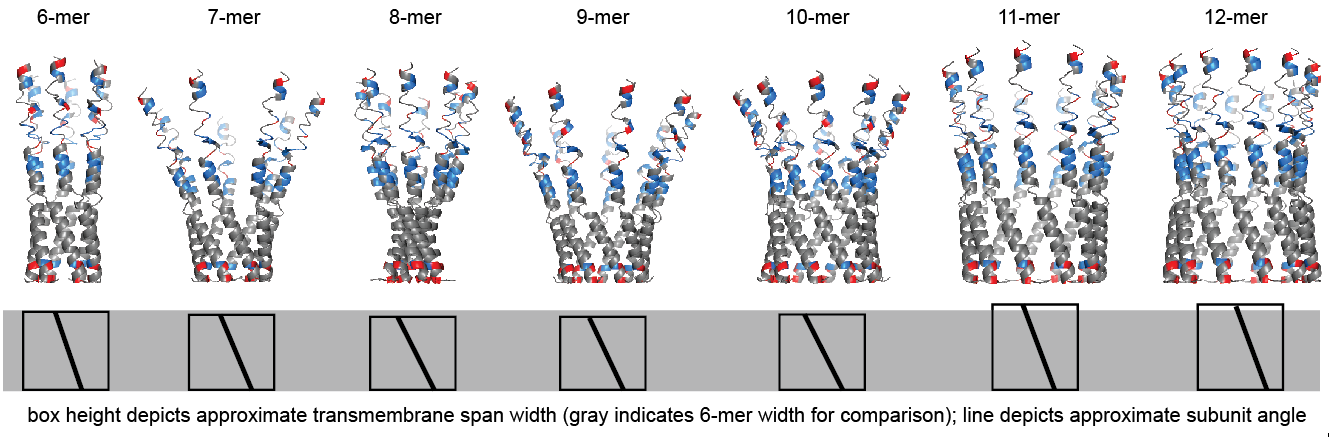

**Figure S3. Hexameric (6-mer) and potentially 11- and 12-mer Mtln complexes have highly interfaced subunits and membrane domain lengths that could more plausibly span mitochondrial membrane widths.**

Possible human Mtln oligomeric complexes generated from i-Tasser singlet model input into GALAXYHomomer are shown with schematic representations depicting subunit angles and transmembrane domain spanning lengths (box heights). Gray shaded box indicates about ~2.8 nm corresponding to the estimated hexameric (6-mer) transmembrane spanning width. Mitochondrial membrane widths are estimated to be ~3 nm.

**Figure S4**

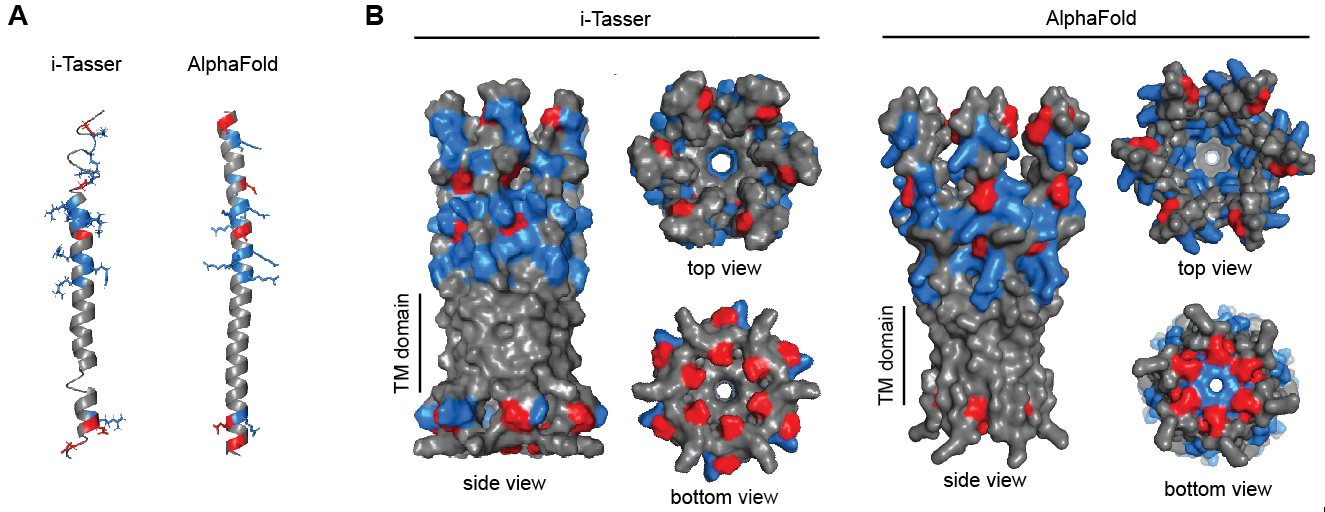

**Figure S4: Mouse Mtln homo-oligomerization may form a hexameric pore-like complex.**

**A.** Singlet models of mouse Mtln generated using default settings on i-Tasser and AlphaFold show linear alpha-helical structures. Models were visualized using PyMOL software. Negatively- and positively-charged amino acids are indicated by red and blue respectively. **B.** Structural models of a mouse Mtln hexamers generated from GALAXYHomomer using singlet i-Tasser and AlphaFold models. Models were visualized using PyMOL. Transmembrane (TM) spanning portions are marked.
